# The snowmelt niche differentiates three microbial life strategies that influence soil nitrogen availability during and after winter

**DOI:** 10.1101/2020.01.10.900621

**Authors:** Patrick O. Sorensen, Harry R. Beller, Markus Bill, Nicholas J. Bouskill, Susan S. Hubbard, Ulas Karaoz, Alexander Polussa, Heidi Steltzer, Shi Wang, Kenneth H. Williams, Yuxin Wu, Eoin L. Brodie

## Abstract

Soil microbial biomass can reach its annual maximum pool size beneath the winter snowpack and is known to decline abruptly following snowmelt in seasonally snow-covered ecosystems. Observed differences in winter versus summer microbial taxonomic composition also suggests that phylogenetically conserved traits may permit winter-versus summer-adapted microorganisms to occupy distinct niches. In this study, we sought to identify archaea, bacteria, and fungi that are associated with the soil microbial bloom overwinter and the subsequent biomass collapse following snowmelt at a high-altitude watershed in central Colorado, USA. Archaea, bacteria, and fungi were categorized into three life strategies (Winter-Adapted, Snowmelt-Specialist, Spring-Adapted) based on changes in abundance during winter, the snowmelt period, and after snowmelt in spring. We calculated indices of phylogenetic relatedness (archaea and bacteria) or assigned functional attributes (fungi) to organisms within life strategies to infer whether phylogenetically conserved traits differentiate Winter-Adapted, Snowmelt-Specialist, and Spring-Adapted groups. We observed that the soil microbial bloom was correlated in time with a pulse of snowmelt infiltration, which commenced 65 days prior to soils becoming snow-free. A pulse of nitrogen (N, as nitrate) occurred after snowmelt, along with a collapse in the microbial biomass pool size, and an increased abundance of nitrifying archaea and bacteria (e.g., Thaumarchaeota, Nitrospirae). Winter- and Spring-Adapted archaea and bacteria were phylogenetically clustered, suggesting that phylogenetically conserved traits allow Winter- and Spring-Adapted archaea and bacteria to occupy distinct niches. In contrast, Snowmelt-Specialist archaea and bacteria were phylogenetically overdispersed, suggesting that the key mechanism(s) of the microbial biomass crash are likely to be density-dependent (e.g., trophic interactions, competitive exclusion) and affect organisms across a broad phylogenetic spectrum. Saprotrophic fungi were the dominant functional group across fungal life strategies, however, ectomycorrhizal fungi experienced a large increase in abundance in spring. If well-coupled plant-mycorrhizal phenology currently buffers ecosystem N losses in spring, then changes in snowmelt timing may alter ecosystem N retention potential. Overall, we observed that the snowmelt separates three distinct soil niches that are occupied by ecologically distinct groups of microorganisms. This ecological differentiation is of biogeochemical importance, particularly with respect to the mobilization of nitrogen during winter, before and after snowmelt.

## Introduction

Snowmelt is a hydrologic event that exerts significant influence on annual water and nitrogen (N) export in seasonally snow-covered, mountainous catchments (Bales et al. 2006). Historically, the timing of snowmelt has coincided with a suite of environmental triggers that characterize the transition from winter to spring, including rising air temperature, longer days having higher photosynthetically active radiation, and greater soil moisture availability (Marks and Dozier 1992, Harpold and Molotch 2015, Musselman et al. 2017). These abiotic cues initiate phenological transitions between winter and spring metabolic states for both plants and soil microbial communities (Richardson et al. 2006, Inouye 2008, Miller-Rushing et al. 2010, Contosta et al. 2016, Ladwig et al. 2016, Thackeray et al. 2016). As a consequence, snowmelt is found to be a critical period that influences nutrient mobilization, assimilation, and retention in terrestrial ecosystems (Brooks et al. 1998, Brooks and Williams 1999, Grogan and Jonasson 2003, Kielland et al. 2006, Campbell et al. 2007). Rising global air temperature has led to reductions in winter snowpack extent, earlier snowmelt dates, and altered growing season length in many mountainous catchments (Mote et al. 2005, Steltzer and Post 2009, Harte et al. 2015, Sloat et al. 2015, Bormann et al. 2018, Prevéy et al. 2019). The ecological consequences of such environmental changes, however, are not well understood (Ernakovich et al. 2014).

Soil microbial biomass production beneath the winter snowpack can be significant, and in some ecosystems, microbial biomass may reach its annual maximum pool size beneath the winter snowpack (Brooks et al. 1998, Lipson et al. 1999, Grogan and Jonasson 2003, Edwards et al. 2006, Larsen et al. 2007, Buckeridge et al. 2010). The microbial bloom in soil during winter immobilizes N within microbial biomass (Brooks et al. 1998). An abrupt collapse in the microbial biomass pool size after snowmelt results in the subsequent release of N from the microbial biomass pool (Lipson et al. 1999, Buckeridge et al. 2010, Isobe et al. 2018). Numerous mechanisms have been hypothesized to induce the microbial biomass crash, including substrate depletion and starvation of the winter community, cell lysis due to soil freeze-thaw events, sudden changes in the osmotic environment with snowmelt, grazing by protozoa and mesofauna, and mortality and replacement of winter-adapted psychrophiles by summer-adapted mesophiles (Jefferies et al. 2010). A better understanding of the factors that promote microbial biomass production under-snow and the crash after snowmelt is necessary because the flux of N released from microbial biomass after snowmelt can be the largest annual pulse of soil N in some ecosystems (Grogan and Jonasson 2003, Schmidt et al. 2007).

In addition to differences in the size of the microbial biomass pool across seasons, bacterial and fungal community composition are also known to differ taxonomically in winter compared to summer (Schadt et al. 2003, Lipson and Schmidt 2004, Aanderud et al. 2013, Isobe et al. 2018), although this has typically been reported at coarse (e.g., phylum) levels of taxonomic resolution. For example, Acidobacteria, Verrucomicrobia, and Bacteriodetes co-exist in alpine soils, but strong successional differences in relative abundance are observed among these phyla during winter and summer (Schmidt et al. 2007). Because two species competing for the same resources cannot persist together indefinitely, such co-occurrence and succession implies that these phyla do not occupy the same niche or partition resources in space or time (Hutchinson 1957). Differences in growth rates and biomass yields among winter and summer bacterial isolates also indicate that phenotypic traits with physiological trade-offs may differentiate winter- and summer-adapted microorganisms (Lipson et al. 2009).

More broadly, an organism’s fitness across varying environmental gradients should be reflected by its abundance in the environment and constrained by a suite of underlying physiological traits, collectively referred to as an organism’s life strategy (Schimel et al. 2007, Placella et al. 2012, Evans et al. 2014, Ho et al. 2017). If such traits are adaptive and phylogenetically conserved (Webb et al. 2002), then quantifying the phylogenetic relatedness of organisms grouped into life strategies may provide insights into the community assembly processes (e.g., niche partitioning) that underlie the microbial bloom beneath the winter snowpack and subsequent biomass collapse after snowmelt.

Within this framework we addressed the following questions: (1) are closely related microorganisms responsible for the overwinter microbial biomass bloom, or is the phenomenon widespread and characteristic among distantly related taxa? (2) what is the mechanism responsible for microbial biomass crash following snowmelt and how widespread is the snowmelt decline among distantly related bacterial, archaeal, and fungal taxa? (3) are there identifiable temporal abundance patterns that differentiate microorganisms into life strategies that can be used to infer an organism’s snowmelt niche?

To address these questions, we sampled soils along a 200-m upland hillslope-to-riparian floodplain transect in a mountainous catchment at the East River Watershed, Colorado, USA (Hubbard et al. 2018). Soil samples were collected over a time-course that began after plant senescence in autumn, through snow accumulation during winter, during snowmelt, and through plant green-up in spring and early summer. We monitored soil temperature and moisture and quantified the dynamics of the soil microbial biomass pool as well as extractable N in soil that was measured as extractable soil nitrate (NO3^-^). Bacterial, archaeal, and fungal species relative abundances were also measured over this seasonal time-course. Here, we show that soil temperature and soil moisture during winter are controlled by winter snowpack dynamics and that snowmelt induces rapid changes in the soil abiotic environment. In addition, snowmelt triggered marked changes in the size of the microbial biomass pool, which determined soil N availability. Finally, we show that specific bacterial, archaeal, and fungal taxa are associated with the rise and fall of microbial biomass overwinter and that the abundance responses can be used to differentiate taxa into three distinct life strategies separated in time by the snowmelt event.

## Materials and Methods

The Upper East River Watershed is located in Gunnison County, CO in the West Elk Mountains near the towns of Crested Butte and Gothic (38°57.5’ N, 106°59.3’ W) and is home to the Rocky Mountain Biological Laboratory. Elevations within the East River watershed range from 2750 to 4000 m. The climate at East River is continental and characterized by periods of persistent snow cover in winter lasting 4 to 6 months (e.g., November through May) followed by an arid summer with intermittent, monsoonal rain events that occur from July through September. The lowest daily air temperatures typically occur in January (−14 ± 3° C), whereas the highest daily air temperatures typically occur in July (23.5 ± 3° C). The average summer air temperature has increased by 0.5 ± 0.1 °C per decade since 1974 (Canberry and Inouye 2014). The average annual precipitation is approximately 1200 cm, with the majority (> 80%) falling as snowfall during winter (Harte and Shaw 1995, Hubbard et al. 2018). The maximum annual snow depth between 1974 to 2017 was 465 cm and the date of snowmelt in spring has advanced by 3.5 ± 2 days per decade since 1974 (Iler et al. 2012).

We sampled a 200-m transect that transitioned from an upland hillslope (hereafter referred to as Hillslope) to a riparian floodplain (hereafter, ‘Floodplain’) adjacent to the main stem of the East River (elevation ∼2760 to 2775 m). We chose this transect because these ecosystem types are representative of dominant land cover adjacent to the East River. Vegetation in the Hillslope is a mix of perennial bunchgrasses (e.g., *Festuca arizonica*), forbs (e.g., *Lupinus* spp., *Potentilla gracilis, Veratrum californicum*), and shrubs (*Artemesia tridentata*), whereas plant communities in the Floodplain are dominated by dwarf birch (*Betula grandulosa*) and mountain willow (*Salix monticola* (Falco et al. 2019)).

Soil temperature, soil volumetric water content, and water potential were measured continuously starting in October 2016 at three locations on the Hillslope and at three locations on the Floodplain. Soil temperature was measured continuously at 6 and 9 cm below the soil surface (sensor model RT-1, 5TE and MPS6, METER). Soil volumetric water content was measured at 9 cm using a time-domain reflectometry probe (model 5TE, METER Group). Soil water potential was measured at 17 cm using MPS6 from METER. We measured snow depth during winter with a marked snow pole or meter stick on the dates of soil sampling that had snow cover at the site.

### Soil Sampling and Field Processing

Soil samples were collected on four dates to characterize the soil microbial biomass carbon (C) pool, extractable soil NO3^-^, and microbial community composition, starting first in autumn after plant senescence (September 12, 2016), at peak winter snow depth (March 7, 2017), during the snowmelt period (May 9, 2017), and following the complete loss of snow and the start of the plant growing season (June 9, 2017). During snow-free times of the year, soils were collected using a 4 cm diameter soil bulk density corer at 12 plots at the Hillslope and at 3 plots at the Floodplain. Soil cores were subsampled and split into three discrete depth increments; 0 to 5 cm, 5 to 15 cm, and 15 cm + below the soil surface.

During periods of winter snow cover (i.e., March and May 2017), snow-pits were dug down to the soil surface at three locations on the Hillslope and one location in the Floodplain in order to sample soils from beneath the snowpack. In each snow-pit, soils were sampled at two adjacent locations separated by more than 1 m using the soil coring method described above. Thus, during the snow-free time of the year, we sampled and analyzed 180 total soil samples (2 ecosystem types x 15 Plots x 2 Time Points x 3 Depths) and 48 soils total during winter (4 Snow-pits x 2 Time Points x 3 Depths x 2 replicate cores). A ∼10 g subsample from each soil core at each depth was placed immediately on dry ice in the field, frozen, and archived for bacterial and fungal community analysis. The remainder of the soil core was allocated to physical and chemical characterization (described below) and was stored at 4 °C until further analysis.

### Soil Physical and Chemical Properties

Gravimetric soil moisture content was measured for each sample on each sampling date by determining the mass of water lost during a 48-hr incubation at 60 °C. Soil pH was determined in a 1:1 w/v slurry in reagent water (18 MΩ resistance; GenPure UltraPure Water System; Thermo Scientific) using an Orion soil pH probe (Thermo Scientific). Total soil organic C and N concentrations and soil δ^13^C and δ^15^N were measured using a Costech ECS 4010 elemental analyzer coupled to a Thermo Delta V Plus isotope ratio mass spectrometer at the Center for Isotope Geochemistry at Lawrence Berkeley National Laboratory (Berkeley, CA).

Extractable nitrate (NO3^-^) as well as microbial biomass carbon (C) were quantified in 0.5M K2SO4 extracts. A 5 g subsample of each soil (field-moist) was extracted in 25 mL of 0.5M K2SO4 on an orbital shaker table for 60 min, then gravity filtered through pre-leached #42 Whatman filter paper and frozen until further analysis. NO3^-^ concentration in the 0.5M K2SO4 extracts was measured colorimetrically using a Versamax microplate spectrophotometer (Molecular Devices) using a modified Greiss reaction protocol (Doane and Horwath 2003). Microbial biomass carbon (C) was measured using the chloroform-fumigation extraction method (Brookes et al. 1985). A 5-g subsample was fumigated with ethanol-free chloroform for 7 days and then extracted as stated above. Microbial biomass C was estimated as organic C measured in the fumigated extract minus organic C measured in the non-fumigated sample (Brookes et al. 1985). We did not apply an extraction correction to account for incomplete microbial biomass lysis during the fumigation. The concentration of dissolved organic C in the fumigated and non-fumigated samples was quantified using the Mn-(III) pyrophosphate oxidation method (Bartlett and Ross 1988). We applied a universal correction factor to the measured organic C concentrations to account for differences in reaction efficiency across soil types (Giasson et al. 2014). All concentrations were corrected for the field-moist water content of the soil and values are reported based on the oven-dried soil mass used for the extractions.

### Illumina library preparation and bioinformatics

Total genomic DNA was extracted in triplicate from each soil sample collected on each date using the DNAEasy Power Soil DNA Extraction Kit (QIAGEN) with the protocol modified to include a 5-min incubation at 65 °C prior to bead-beating to increase lysis efficiency. Replicate soil DNA extracts were combined prior to PCR amplification. PCR reaction conditions are summarized in the Supplemental Methods. Purified PCR products were pooled in equimolar concentrations and sequenced on a single lane for 300-bp paired-end v3 Illumina MiSeq sequencing conducted at the Vincent J. Coates Genomics Sequencing Laboratory at the University of California, Berkeley.

Forward and reverse reads were aligned and paired using usearch (v8.1.1861, (Edgar 2010)) *fastq_mergepairs* command (maximum diff = 3). The aligned reads were quality filtered (command *fastq_filter* with *-fastq_trunclen=230, -fastq_maxee=0.1*), concatenated into a single fasta file, and singletons were removed (command *sortbysize* with *minsize=2*). These filtered sequences were used for operational taxonomic unit (OTU) clustering with the *uparse* pipeline (Edgar 2013) setting the OTU cut-off threshold to 97%. Chimeric sequences were filtered with uchime (Edgar et al. 2011) with reference to the ChimeraSlayer database downloaded from http://drive5.com/uchime/gold.fa. OTU abundances across individual samples were calculated by mapping chimera-filtered OTUs against the quality-filtered reads (command *usearch_global* with *-strand plus -id 0.97*). Taxonomy was assigned to each OTU by a Naïve Bayes classifier using the assign_taxonomy.py command in QIIME (Wang et al. 2007) with reference to the SILVA database accessed from mothur (Schloss et al. 2009) release 119: https://mothur.org/wiki/Silva_reference_files#Release_119). For phylogenetic inference of bacterial and archaeal OTUs, representative sequences for each bacterial OTU were aligned to a SILVA SEED sliced alignment using the PyNAST algorithm (Caporaso et al. 2010) and archaeal and bacterial phylogeny was inferred using FastTree (Price et al. 2010).

This workflow resulted in 18,716 archaeal and bacterial OTUs and 4,194 fungal OTUs. OTUs were further filtered to include only OTUs that occurred in at least 25% of samples across sampling dates. Fungal OTUs were also assigned to functional guilds by referencing annotated databases using the open-source software FUNGuild (Nguyen et al. 2016). FUNGuild assigned functional attributes (e.g., trophic mode, guild) to 1528 fungal OTUs. Fungal OTUs with ‘unknown’ FUNGuild functional annotations were also included in subsequent downstream analyses. Raw de-multiplexed sequences have been archived in the NCBI Bioproject database under accession no. PRJNA587134.

### Statistical Analysis

All statistical analyses were completed using R v 3.5.2 (R Development Core Team 2010). Differences in soil chemical properties between locations (Hillslope versus Floodplain) were tested using linear-mixed effect models using the R package ‘nlme’ (Pinheiro et al. 2016). *Location* was the fixed-effect and *Plot* was the random-effect in the model. The effect of sampling date on the size of the microbial biomass C pool, extractable soil NO3^-^, and gravimetric water content were also assessed using linear-mixed effect models with *Time of Sampling* as the fixed-effect and *Plot* as the random-effect in the model. Differences in bacterial and fungal community composition across sampling dates was determined by permutational multivariate analysis of variance (perMANOVA, permutations *n* = 999) using the R package ‘vegan’ (Oksanen et al. 2011). Dissimilarity between samples was calculated as Bray-Curtis distances for fungi or weighted-unifrac distances for bacteria and archaea (Bray and Curtis 1957, Lozupone et al. 2007).

We identified OTUs that had a statistically significant change in abundance between at least one of three paired time periods [September to March (i.e., ‘Winter’), March to May (i.e., ‘Snowmelt’), May to June (i.e., ‘Spring’)] by calculating log2foldchanges in relative abundance between time-points using the R package ‘gtools’ (Warnes et al. 2018). A 95% confidence interval for the log2foldchange was derived by applying a formula for error propagation for the product of a ratio,

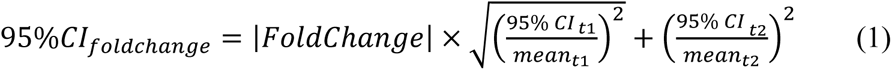

where meant1,t2 was the mean relative abundance of each OTU across sampling locations on each date and 95% CI t1,t2 was the 95% confidence interval of each OTU on each date. P-values were calculated from the confidence intervals (Altman and Bland 2011) and were adjusted for multiple comparisons by the Benjimini and Hochberg false discovery rate (‘fdr’) method using the R package ‘stats’. Only OTUs with at least one significant (e.g., FDR-adjusted *P* < 0.05) log2foldchange across time points were included in further analysis (4591 bacterial OTUs and 1110 fungal OTUs).

Next, we applied agglomerative hierarchical clustering to group OTUs according to differences in relative abundance patterns across sampling time points (e.g. Placella et al. 2012, Evans et al. 2014). Distances between clusters were calculated by the average-linkage method. Dendrograms were visualized using ‘ggdendro’ (de Vries and Ripley 2016) and cut at even height to categorize OTUs into one of three life strategies. Here we define life strategy to be an assortment of physiological traits, selected for by abiotic and biotic factors, that determine an organism’s acquisition of resources, growth and reproduction, and responses to varying environmental gradients (Chagnon et al. 2013, Ho et al. 2017, Malik et al. 2018). In our study, we assume that changes in relative abundance underlie distinct life strategies related to winter snowpack dynamics and reflect each organism’s degree of adaptation and fitness during winter, snowmelt, and spring.

Lastly, we calculated two measures of phylogenetic community relatedness within life strategies for bacteria and archaea, the net relatedness index (NRI) and the nearest taxon index (NTI), using the R package ‘picante’ with a null model based on random taxa shuffling and 1000 permutations (Webb et al. 2002, Kembel et al. 2010). Phylogenetic clustering (positive NRI/NTI) indicates that taxa in a group are more phylogenetically related than expected, compared to a random sampling from the regional species pool. Phylogenetic clustering suggests a group of taxa that are ecologically similar and share a common niche with traits that have been retained through speciation events (i.e., phylogenetic niche conservatism, Crisp and Cook 2012). On the other hand, phylogenetic overdispersion (negative NRI/NTI) indicates a group of organisms less phylogenetically related than expected, as compared to the regional species pool. Phylogenetic overdispersion can arise from community assembly processes (e.g., trophic interactions, competitive exclusion) that result in a group of ecologically dissimilar taxa with non-overlapping niches (i.e., niche partitioning) or adaptive traits with a broad phylogenetic distribution.

## Results

### Soil characteristics and winter snowpack control over soil temperature and moisture

Average gravimetric soil moisture content and total soil C were greater on the Floodplain compared to the Hillslope (Table 1). Microbial biomass C in the top-soil (0 to 5 cm depth) was also approximately three times greater on the Floodplain compared to Hillslope, but was comparable between the Hillslope and Floodplain in soils deeper than 5 cm (Table 1). Across all sampling depths, soils on the Hillslope had consistently greater δ^13^C and δ^15^N values than soils in the Floodplain (Table 1).

**Table 1.** Soil properties measured in the Hillslope and Floodplain watershed locations. Differences in means among Locations are denoted with different letters and represent statistically significant differences (Tukey post-hoc *P* ≤ 0.05).

| Depth |  | Hillslope |  | Floodplain |  |
| --- | --- | --- | --- | --- | --- |
|  |  | Mean | Std Error | Mean | Std Error |
| 0 to 5 cm | Soil Moisture | 0.42<sup>b</sup> | 0.10 | 1.16<sup>a</sup> | 0.3 |
|  | Soil pH | 6.7 | 0.1 | 6.8 | 0.2 |
|  | Microbial Biomass C | 203.7<sup>b</sup> | 21.6 | 629.4<sup>a</sup> | 117.5 |
|  | Extractable Organic C | 99.4 | 10.5 | 101.2 | 29.3 |
|  | Total Soil organic C | 4.2<sup>b</sup> | 0.3 | 7.4<sup>a</sup> | 2.1 |
|  | Total Soil organic N | 0.42 | 0.01 | 0.58 | 0.14 |
| | Soil $\delta^{13}\text{C}_{\text{VPDB}}$ | -24.3<sup>b</sup> | 0.1 | -26.6<sup>a</sup> | 0.5 |
| | Soil $\delta^{15}\text{N}_{\text{AIR}}$ | 4.0<sup>a</sup> | 0.1 | 2.6<sup>b</sup> | 0.9 |
| 5 to 15 cm | Soil Moisture | 0.34 | 0.1 | 0.72 | 0.1 |
|  | Soil pH | 6.7 | 0.1 | 6.9 | 0.1 |
|  | Microbial Biomass C | 217.2 | 24.8 | 276.8 | 35.7 |
|  | Extractable Organic C | 85.3 | 15.6 | 80.3 | 25.9 |
|  | Total Soil organic C | 3.3<sup>b</sup> | 0.2 | 7.0<sup>a</sup> | 0.4 |
|  | Total Soil organic N | 0.35<sup>b</sup> | 0.01 | 0.53<sup>a</sup> | 0.02 |
| | Soil $\delta^{13}\text{C}_{\text{VPDB}}$ | -24.3<sup>a</sup> | 0.1 | -26.4<sup>b</sup> | 0.2 |
| | Soil $\delta^{15}\text{N}_{\text{AIR}}$ | 4.0<sup>a</sup> | 0.2 | 2.0<sup>b</sup> | 1.2 |
| 15 cm + | Soil Moisture | 0.32 | 0.07 | 0.84 | 0.2 |
|  | Soil pH | 6.6 | 0.1 | 7.0 | 0.1 |
|  | Microbial Biomass C | 170.5 | 18.2 | 158.3 | 35.4 |
|  | Extractable Organic C | 61.7 | 7.0 | 127.2 | 35.0 |
|  | Total Soil organic C | 3.2 | 0.2 | 4.7 | 1.4 |
|  | Total Soil organic N | 0.34 | 0.01 | 0.57 | 0.08 |
| | Soil $\delta^{13}\text{C}_{\text{VPDB}}$ | -24.0<sup>a</sup> | 0.1 | -25.6<sup>b</sup> | 0.5 |
| | Soil $\delta^{15}\text{N}_{\text{AIR}}$ | 4.4<sup>a</sup> | 0.2 | 1.7<sup>b</sup> | 0.3 |

(Units – Soil moisture ( $\text{g H}_2\text{O}^{-1} \text{ g soil}^{-1}$ ), microbial biomass C and extractable organic C ( $\mu\text{g C g soil}^{-1}$ ), total soil C & N (weight %), soil $\delta^{13}\text{C}$ and $\delta^{15}\text{N}$ are the isotopic composition of carbon and nitrogen in the conventional $\delta$ -notation (per mil, ‰) relative to Vienna Pee Dee Belemnite (VPDB) for carbon and relative to air for nitrogen.

The onset of winter snowpack accumulation, persistence of snow cover, and complete loss of snow (approximately May 15, 2017) resulted in several different soil temperature and moisture regimes between November 2016 and July 2017 (Table 2, Figure 1). Soil temperature at 6 cm depth was less than 0 °C on the Hillslope in early December prior to the development of a persistent snowpack (Figure 1a). Snow accumulation and persistence coincided with a gradual increase in soil temperature to slightly above 0 °C from January throughout the remainder of winter. Complete loss of snow led to higher soil temperatures at both the Hillslope and Floodplain (Figure 1a; Table 2) and trends in soil temperature in spring generally tracked trends in air temperature after snowmelt. We did not observe soil freeze-thaw cycles during or after snowmelt, but did observe soil freeze-thaw cycles in the winter time-period (Supplemental Table 1).

**Figure 1.**
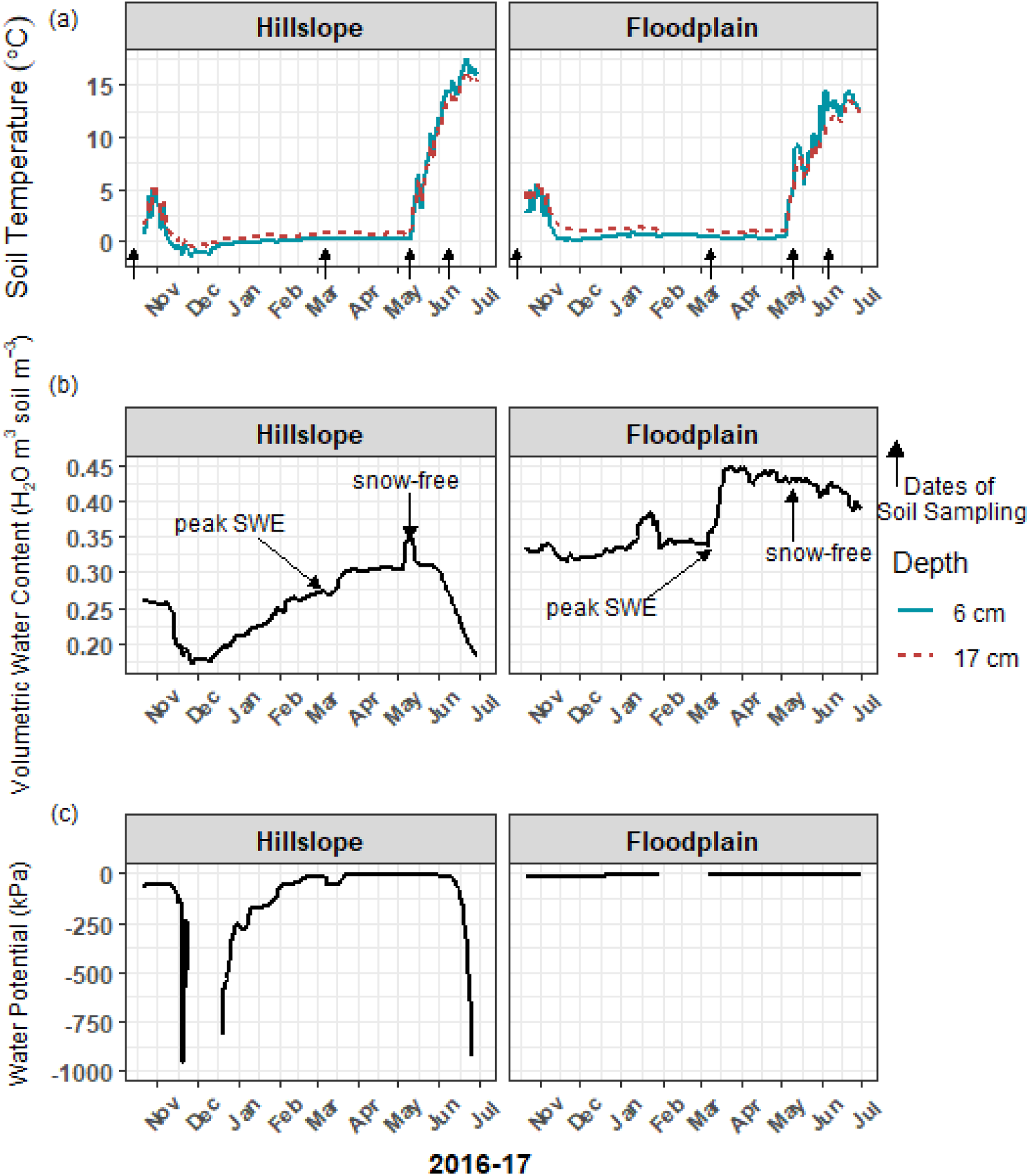
Soil temperature (a) at 6 cm and 17 cm below the soil surface remained above 0 °C when soils were covered with snow during winter. Loss of snow cover in May 2017 resulted in a rapid increase in soil temperature. The onset of snowmelt in March triggered a large increase in soil volumetric water content (b), as well as soil water potential (c), which lasted through early June 2017. Volumetric water content was measured at 9-cm depth and water potential was measured at 17-cm depth below the soil surface. Arrows indicate the dates of soil sampling.

**Table 2.** Snowpack effects on soil temperature and moisture. Time periods were binned based on snow accumulation and loss as well as regime shifts in temperature and moisture. Winter was operationally defined as 10/21/2016 – 03/07/2017, Snowmelt was 03/08/2017-05/07/2017, and Spring was 05/08/2017-6/30/2017. Means and standard errors were calculated from daily means within a binned time period. Snow depth data were collected at multiple locations within a site (Hillslope and Floodplain) on the dates of soil sampling only. Significant differences in means among time-periods are denoted with different letters (*P* ≤ 0.05).

| Depth |  | Winter |  | Snowmelt |  | Spring |  |
| --- | --- | --- | --- | --- | --- | --- | --- |
|  |  | Mean | Std Error | Mean | Std Error | Mean | Std Error |
| Hillslope | Snow Depth | 144.1 | 4.1 | 15.4 | 3.1 | 0 | 0 |
|  | Soil T (6 cm) | 0.2<sup>c</sup> | 0.1 | 0.3<sup>b,c</sup> | 0.1 | 11.1<sup>a</sup> | 0.4 |
|  | Soil T (17 cm) | 0.7<sup>c</sup> | 0.1 | 0.7<sup>b,c</sup> | 0.1 | 10.4<sup>a</sup> | 0.4 |
|  | VWC | 0.22<sup>c</sup> | 0.01 | 0.30<sup>a</sup> | 0.01 | 0.27<sup>b</sup> | 0.01 |
|  | Water | - |  | - |  | - |  |
|  | Potential | 721.0<sup>c</sup> | 85.6 | 19.0<sup>a,b</sup> | 1.2 | 197.0<sup>b</sup> | 40.2 |
| Floodplain | Snow Depth | 164.7 | 1.2 | 8.6 | 2.4 | 0 | 0 |
|  | Soil T (6 cm) | 1.0<sup>b</sup> | 0.1 | 0.5<sup>c</sup> | 0.1 | 10.8<sup>a</sup> | 0.3 |
|  | Soil T (17 cm) | 1.9<sup>c</sup> | 0.1 | 1.0<sup>b</sup> | 0.1 | 9.6<sup>a</sup> | 0.3 |
|  | VWC | 0.33<sup>c</sup> | 0.01 | 0.43<sup>a,b</sup> | 0.01 | 0.42<sup>a</sup> | 0.01 |
|  | Water |  |  |  |  |  |  |
|  | Potential | -12.1<sup>c</sup> | 0.1 | -8.8<sup>a,b</sup> | 0.1 | -8.8<sup>a</sup> | 0.1 |

(Units – snow depth (cm), temperature (°C), VWC-volumetric water content ( $\text{m}^3 \text{H}_2\text{O soil m}^{-3}$ ), water potential (kPa))

Soils in winter generally had lower volumetric water content and lower soil water potential compared to soils collected during snowmelt or spring (Table 2). Because over 90% of the winter snowpack was melted between early March and early May 2017, we refer to that time-period as the snowmelt period (Table 2). Snowmelt resulted in an increase in soil volumetric water content and soil water potential (Figure 1b,c; Table 2) with an extended period of soil saturation beginning more than 60 days prior to the complete loss of winter snow. At the Hillslope, soil moisture declined rapidly after snowmelt as plants broke winter dormancy in spring (Figure 1b,c). In contrast, soils on the Floodplain remained saturated long after snowmelt (Figure 1c; Table 2). In summary, the winter period was characterized generally by deep snow with cold and dry soils, the snowmelt period by the rapid loss of the winter snowpack with cold and wet soils, and the spring soil environment was characterized by rapid warming and drying (Table 2, Figure 1a,b,c).

### Microbial biomass bloom and crash with subsequent pulse of soil N

Soil microbial biomass and the extractable soil nitrate (NO3^-^) pool responded strongly to variations in snowpack depth, soil temperature, and moisture (Figure 2a,b,c; Figure 1). For example, soil microbial biomass C increased 2-to 5-fold at all three soil depths (0 to 5 cm, 5 to 15 cm, 15 cm +) during the 65-day snowmelt period on the Hillslope (e.g., March to May, Figure 2b) and trends in microbial biomass pool size generally tracked trends in soil water content (Figure 2a). Similar to observations for the Hillslope, microbial biomass C in the Floodplain also increased during snowmelt in the shallowest soils (e.g., 0 to 5 cm) (Figure 2b), however, this was not observed in soils sampled more than 5 cm below the soil surface. Soil microbial biomass decreased dramatically after snowmelt (between May and June) on the Hillslope (Figure 2b), which coincided with a substantial pulse of extractable soil NO3^-^ (Figure 2c). Although microbial biomass also collapsed on the Floodplain in topsoils after snowmelt (Figure 2b), in contrast to the Hillslope, we did not observe an increase in the soil NO3^-^ pool size after snowmelt in the Floodplain (Figure 2c).

**Figure 2.**
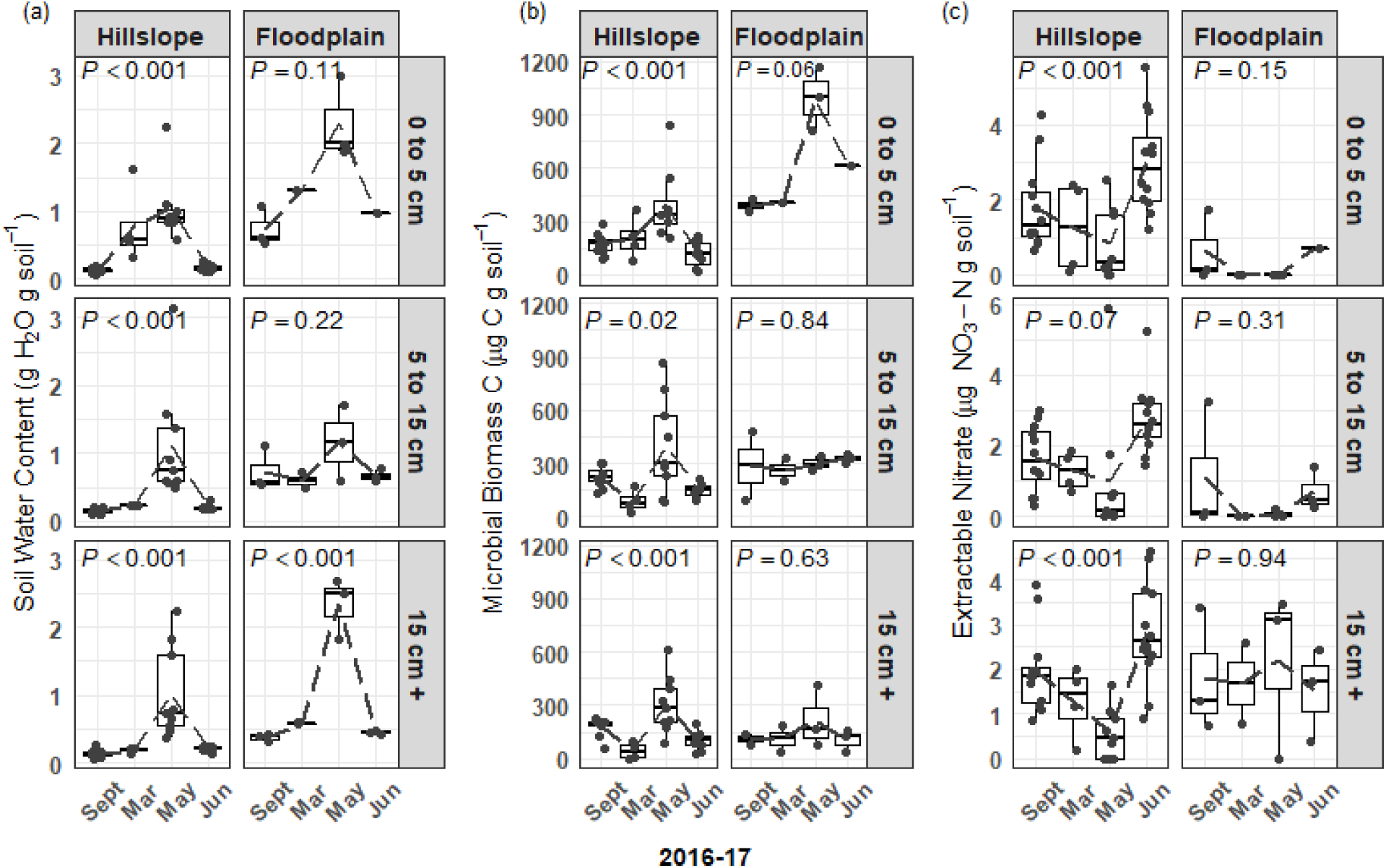
Soil water content (a) and soil microbial biomass (b) in the Hillslope and Floodplain. A pulse of extractable soil nitrate (c) was observed in the Hillslope after snowmelt in June 2017. P-values are the outcome of mixed-models testing for the effect of time of sampling on soil water content, microbial biomass, and extractable nitrate.

Similar to changes in the sizes of soil microbial biomass pools, bacterial and fungal community structure varied significantly during winter, snowmelt, and following the loss of snow in spring (Figure 3a-d). Date of sampling was a significant factor explaining bacterial community structure in both the Hillslope (perMANOVA pseudo-R^2^ = 0.11, *P* ≤ 0.001) and Floodplain (perMANOVA pseudo-R^2^ = 0.33, *P* ≤ 0.001) (Figure 3a,b). Fungal community structure was likewise significantly related to the date of soil sampling in both the Hillslope (pseudo-R^2^ = 0.05, *P* ≤ 0.001) and Floodplain (perMANOVA pseudo-R^2^ = 0.22, *P* < 0.001) (Figure 3c,d). Depth was a significant independent factor for both bacteria (perMANOVA pseudo-R^2^ = 0.33, *P* < 0.001) and fungi (perMANOVA pseudo-R^2^ = 0.05, *P* < 0.001) in the Hillslope only, but not in the Floodplain. We did not observe a significant Date x Depth of sampling interaction for either bacteria or fungi in either the Hillslope or Floodplain (perMANOVA *P* ≥ 0.05 in all cases).

**Figure 3.**
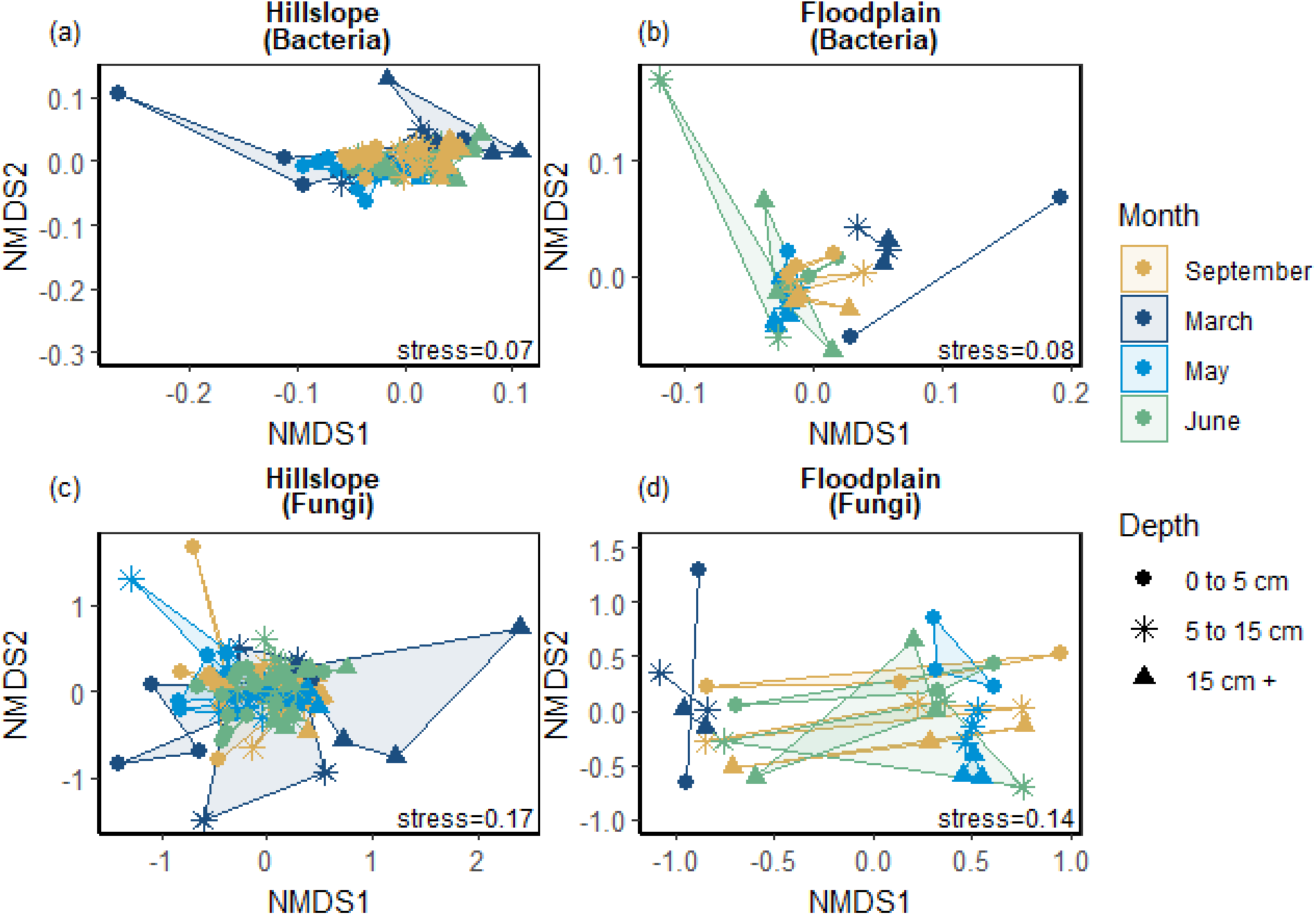
Bacterial community structure represented by non-metric dimensional scaling (NMDS) in the Hillslope (a) and Floodplain (b) as well as fungal community structure in the Hillslope (c) and Floodplain (d).

### Snowmelt selected for phylogenetically clustered bacterial life strategies

Bacterial and archaeal species were grouped together into one of three life strategies based on changes in relative abundance in response to snow accumulation, snowmelt, and the onset of spring (Figure 4a-h). As a group, Winter-Adapted archaea and bacteria had the highest group relative abundance (i.e., all Winter-Adapted OTUs in group summed together) in March (Figure 4g, h), a time when winter snow depth was greatest and soils were relatively cool and dry (Table 2). The relative abundance of Snowmelt-Specialist archaea and bacteria increased 1.8 to 2.4-fold on the Hillslope (Figure 4g) and 2-to 6-fold in the Floodplain (Figure 4h) during the snowmelt period (i.e., March – May). The majority of bacterial and archaeal taxa across life strategies were Spring-Adapted (Table 3). Spring-Adapted archaea and bacteria group relative abundance increased after snowmelt and ranged from 20 to 45% of the total community in the Hillslope or 5 to 8% of the total community in the Floodplain (Figure 4g, h).

**Figure 4.**
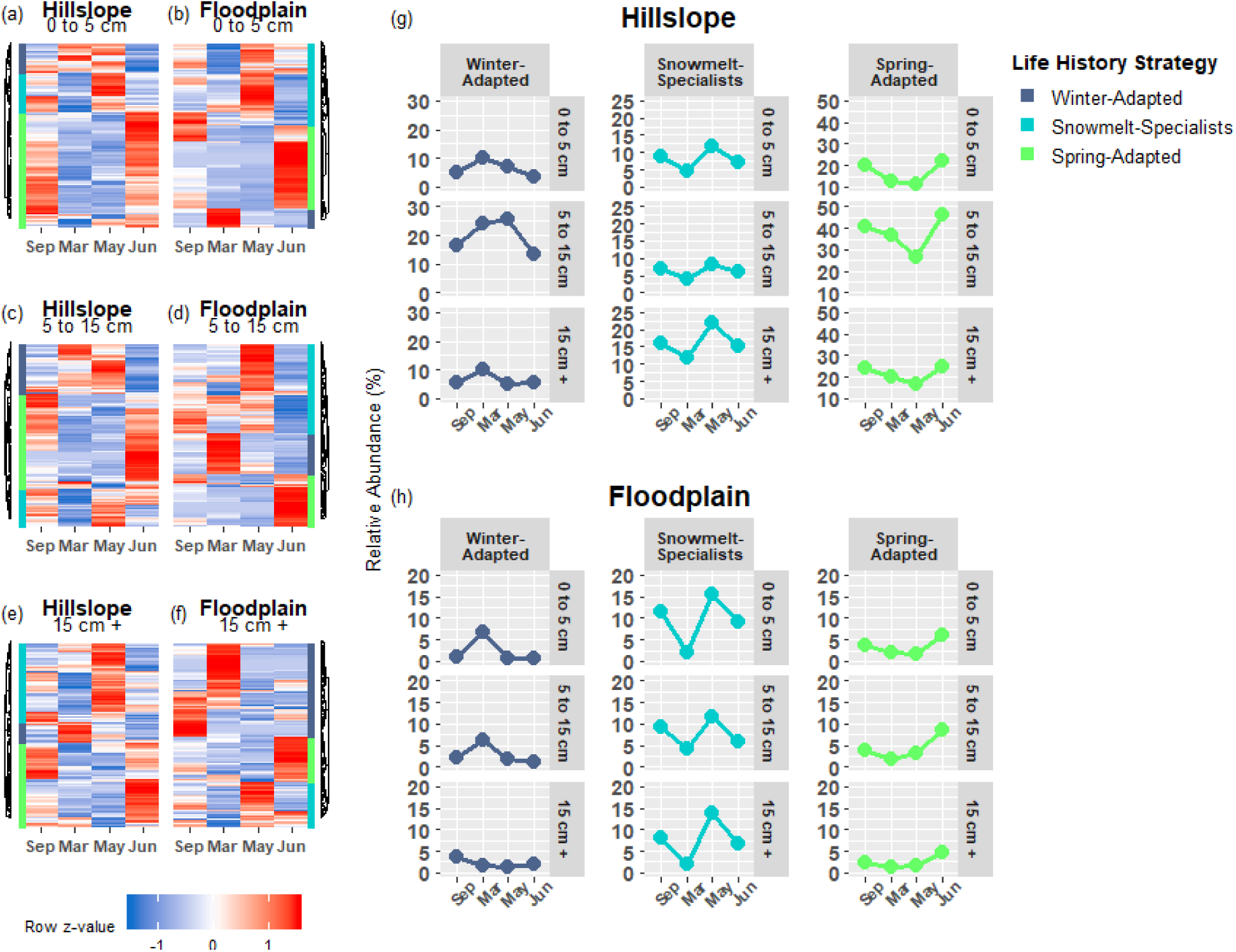
Archaeal and bacterial OTUs that had a significant change in abundance between any two sampling time points (September, March, May, June) were grouped by hierarchical clustering in the Hillslope (5a,c,e) and Floodplain (5b,d,f). Three life-history strategies related to winter snow cover, snowmelt, and loss of snow cover were identified and the group (i.e. sum of all OTUs in group) relative abundance patterns of each strategies are shown in panels (g,h)

**Table 3.** Phylogenetic relatedness of bacteria and archaea grouped by life-history strategy. Values in bold indicate 0.025 > *P* > 0.975. Positive NTI/NRI values indicate that groups are more phylogenetically related (clustered) than expected by chance, whereas negative values indicate that groups are less phylogenetically related (overdispersed) than expected by chance.

| Location | Depth | Life History Strategy | #OTUs | NRI | NTI |
| --- | --- | --- | --- | --- | --- |
| Hillslope | 0 to 5 cm | Winter-Adapted | 55 | -0.48 | **2.18** |
|  |  | Snowmelt-Specialists | 335 | **-2.15** | 1.13 |
|  |  | Spring-Adapted | 1093 | **5.28** | **2.75** |
|  | 5-15 cm | Winter-Adapted | 138 | -1.67 | **5.16** |
|  |  | Snowmelt-Specialists | 433 | -0.89 | 0.33 |
|  |  | Spring-Adapted | 1120 | **9.53** | **5.21** |
|  | 15 cm + | Winter-Adapted | 132 | **1.69** | **3.37** |
|  |  | Snowmelt-Specialists | 461 | -0.16 | 0.31 |
|  |  | Spring-Adapted | 771 | **1.77** | 0.47 |
| Floodplain | 0 to 5 cm | Winter-Adapted | 83 | **2.05** | **3.49** |
|  |  | Snowmelt-Specialists | 269 | **3.13** | **2.25** |
|  |  | Spring-Adapted | 631 | 0.05 | 0.19 |
|  | 5-15 cm | Winter-Adapted | 123 | -0.24 | **3.32** |
|  |  | Snowmelt-Specialists | 153 | **3.48** | **3.88** |
|  |  | Spring-Adapted | 387 | **3.63** | -0.01 |
|  | 15 cm + | Winter-Adapted | 119 | 1.60 | -1.27 |
|  |  | Snowmelt-Specialists | 200 | **3.49** | **3.07** |
|  |  | Spring-Adapted | 276 | 1.32 | 1.00 |

Generally speaking, the direction of the phylogenetic relatedness patterns within archaeal and bacterial life strategies were most often towards phylogenetic clustering (positive NRI/NTI values), however, the responses were not always observed at every sampling depth in the Hillslope or Floodplain (Table 3). For example, Spring-Adapted archaea and bacteria were phylogenetically clustered across all soil depths in the Hillslope based on the Net Relatedness Index (NRI, Table 3). In the Floodplain, Snowmelt-Specialist archaea and bacteria were similarly phylogenetically clustered across all sampling depths. In contrast to observations for the Floodplain, Snowmelt-Specialist archaea and bacteria sampled 0 to 5 cm below the soil surface were phylogenetically overdispersed on the Hillslope (NRI = -2.15, Table 3).

*Bradyrhizobium* (α-Proteobacteria)*, Tardiphaga* (α-Proteobacteria)*, Sphingomonas* (α-Proteobacteria), and *Massilia* (β-Proteobacteria) collectively accounted for 73% of the total increase in group relative abundance of Winter-Adapted bacteria at the Hillslope during winter (i.e., September 2016 to March 2017; Figure 5a). In the Floodplain, the DA101 soil group (Verrucomicrobia), *Thaumarchaeota* spp., and *Pedobacter* (Bacteroidetes) likewise contributed substantially to the increase in relative abundance of Winter-Adapted bacteria (Figure 5b). *Streptomyces* (Actinobacteria) and *Candidatus* Nitrososphaera (Thaumarchaeota) together accounted for 11% of the total increase in relative abundance of Snowmelt-Specialist archaea and bacteria from March to May on the Hillslope (Figure 5c), whereas Bacteriovoraceae (δ-Proteobacteria; 44% contribution alone) dominated the response of Snowmelt-Specialists in the Floodplain. Various types of Acidobacteria [e.g., Acidobacteriaceae (Subgroup 1) spp., Subgroup 2 spp., *Candidatus* Solibacter, RB41 spp.] composed the largest fraction of Spring-Adapted bacteria in the Hillslope (Figure 5e). Nitrifying taxa (e.g. *Nitrospira* spp. and *Thaumarchaeota* spp.) also contributed 7% to the increase of Spring-Adapted archaea and bacteria from May to June on the Hillslope (Figure 5e). Similar to the Hillslope, other potentially nitrifying organisms belonging to the phylum Nitrospirae (e.g., *4-29* spp.) were major contributors to the increase of Spring-Adapted bacteria after snowmelt and during plant-green up in the Floodplain (Figure 5f). All bacterial OTUs with life strategies assigned are listed in Supplemental Table 2.

**Figure 5.**
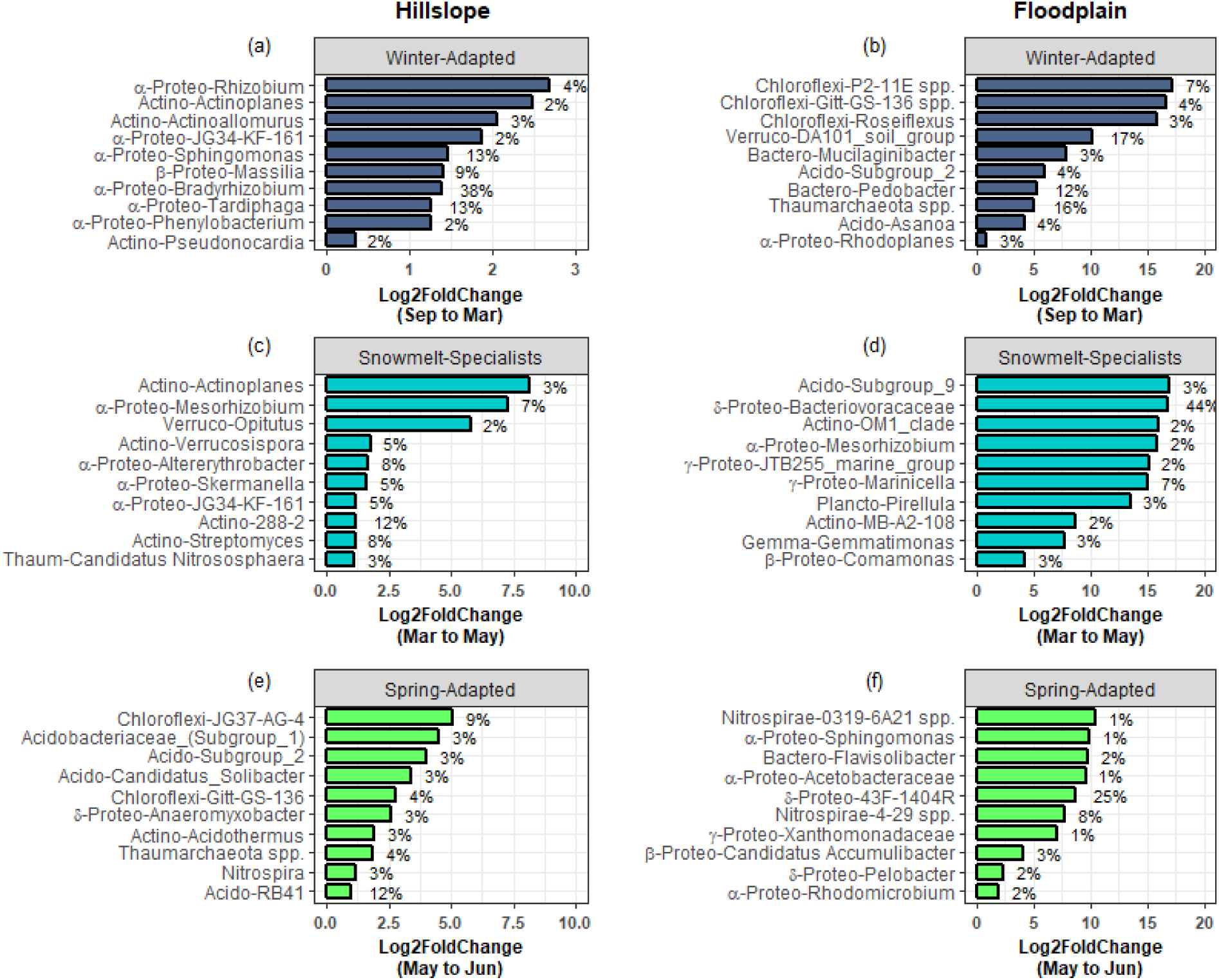
The top 10 archaeal and bacterial phylotypes within each life-history strategy. Phylotypes were ranked by their percent change in abundance between the specified sampling time-points (*x*-axis). The bars represent log2 fold changes and the percent contributions to the change in group relative abundance are given at the side of the bar. Abbreviations for taxonomy (phylum or subphylum level) are Acido – Acidobacteria; Actino – Actinobacteria; Bactero – Bacteriodetes; Gemmo – Gemmatimonadetes; Plancto - Planctomycetes ;α-Proteo – Alphaproteobacteria; β-Proteo – Betaproteobacteria ; δ-Proteo – Deltaproteobacteria; γ-Proteo – Gammproteobacteria; Verruco - Verrucomicrobia

### Fungal taxa and functional groups were also differentiated by snowmelt niche

Winter-Adapted, Snowmelt-Specialist, and Spring-Adapted fungi were also observed across all three soil depths in both the Hillslope and Floodplain (Figure 6a-f). Similar to the archaeal and bacterial communities, Winter-Adapted fungi had the highest group abundance (∼8% of total fungal community) at peak winter snow depth (Figure 6g). Snowmelt-Specialist fungi increased in abundance during the snowmelt period (i.e., March to May), and Spring-Adapted fungi had the highest group abundance during plant green-up in June (Figure 6g,h). The majority of fungal taxa in the Floodplain were Spring-Adapted, with far fewer fungal species categorized as either Winter-Adapted or Snowmelt-Specialists (Figure 6b,d,f, Table 4). For example, Spring-Adapted fungi accounted for 27 to 43% of the total fungal community in the Floodplain in June (Figure 6h), whereas Winter-Adapted and Snowmelt-Specialist fungi were rarely more more than 2% of the total fungal community at any time.

**Figure 6.**
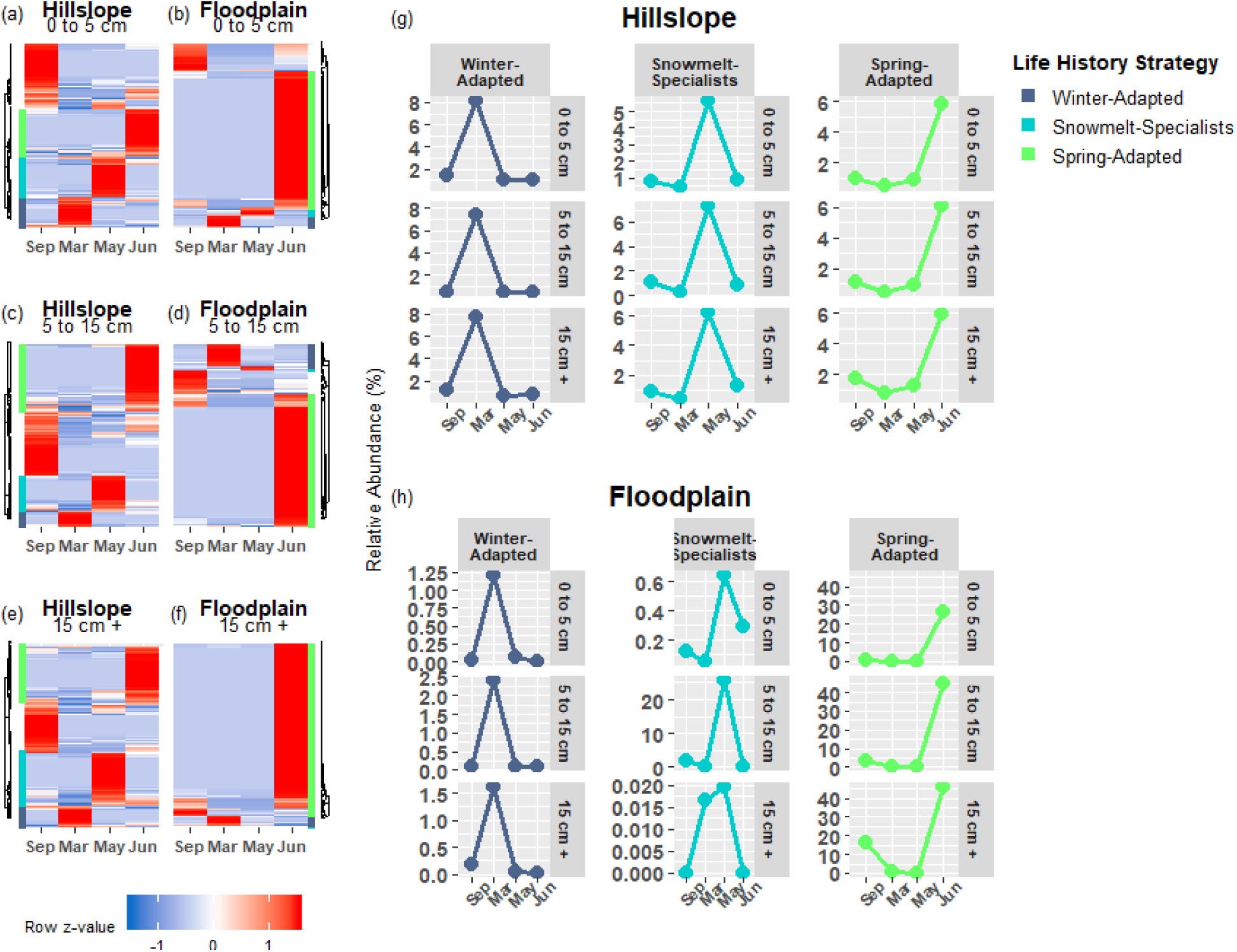
Fungal OTUs that had a significant change in abundance between any two sampling time points were grouped by hierarchical clustering in the Hillslope (a,c,e) and Floodplain (b,d,f). Similar to bacterial OTUs, we observed Winter-Adapted, Snowmelt-Specialist, and Spring-Adapted fungi across all depths and in both the Hillslope and Floodplain.

**Table 4.**
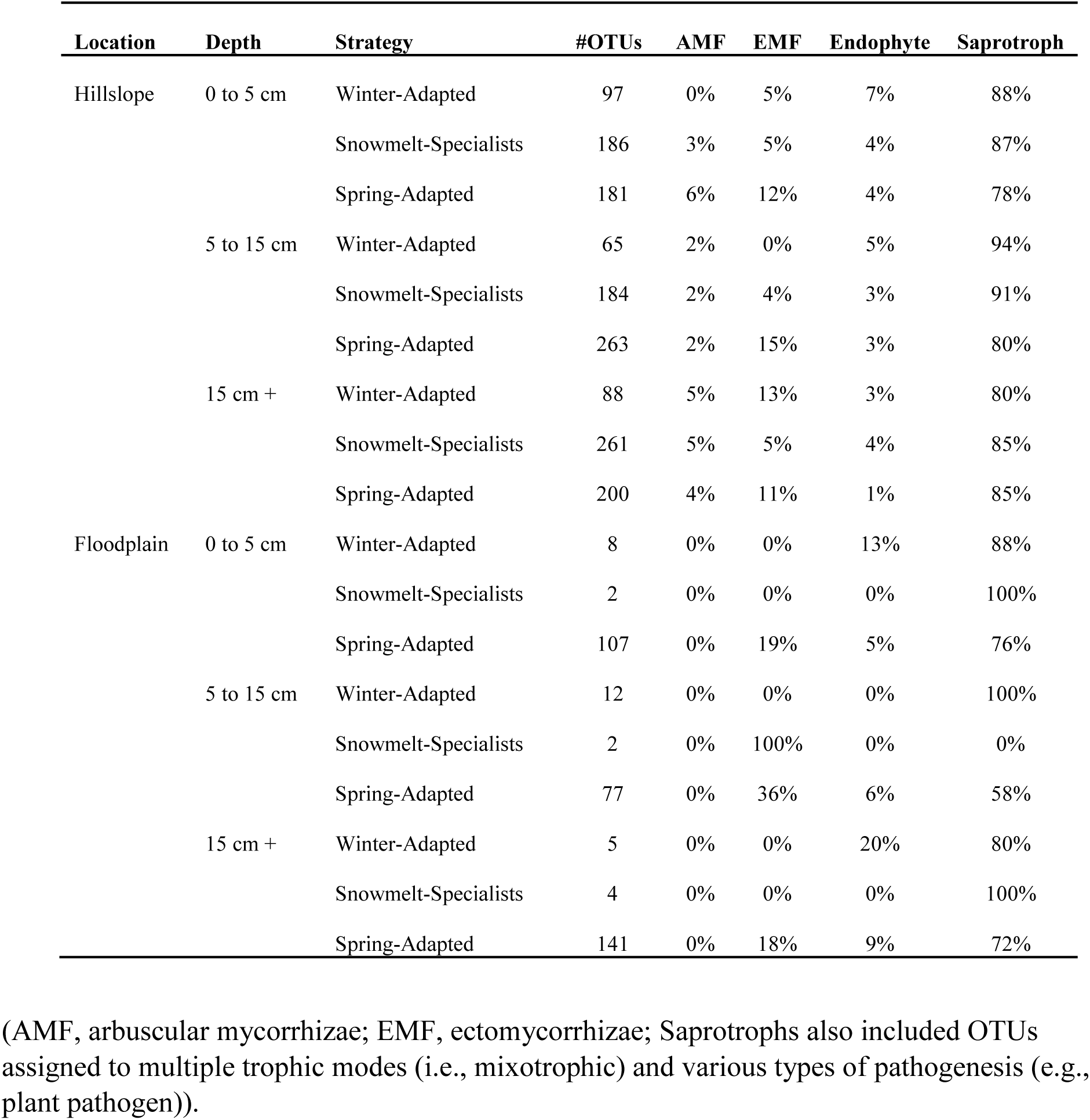
Composition of functional guilds for fungal life-history strategies related to snowmelt. The percent within a functional group was calculated by dividing the total number of OTUs within the functional group by the total numbers of OTUs within a life-history strategy at each depth.

Saprotrophy was the dominant fungal trophic mode at both the Hillslope and Floodplain (Table 4). For example, across all soil depths, 80 to 100% of Winter-Adapted fungal taxa were classified as saprotrophic. Symbiotic fungi that associate with plant roots (e.g., arbuscular mycorrhizae, ectomycorrhizae, and root-endophytes) had higher prevalence within the Spring-Adapted life strategy compared to Winter-Adapted or Snowmelt-Specialist fungi (Table 4). Across soil depths, root-associated fungi constituted 16 to 22% of all Spring-Adapted fungi on the Hillslope and 24 to 38% of Spring-Adapted fungi in the Floodplain. Ectomycorrhizal fungi were the predominant root-associated functional group, especially in the Floodplain, where arbuscular mycorrhizal fungi were not observed among any of the life strategies (Table 4).

*Genabea* (Pezizomycetes), *Psilocybe* (Agaricomycetes), and *Crepidotus* (Agaricomycetes) collectively accounted for 77% of the increase in group relative abundance observed for Winter-Adapted fungi occurring from September to March on the Hillslope (Figure 7a). A few select fungal genera also contributed disproportionately to changes in group abundance for Snowmelt-Specialists and Spring-Adapted fungi during and after snowmelt (Figure 7c,d,e,f). For example, *Pterula* (Agaricomycetes) alone accounted for 53% of the increase in relative abundance for Snowmelt-Specialist fungi during the snowmelt period (i.e., March to May) and *Tricholoma* (Agaricomyetes), *Botrytis* (Leotiomycetes), and *Cuphophyllus* (Agaricomycetes) together accounted for 48% of the increase in group abundance for Spring-Adapted fungi from May to June on the Hillslope (Figure 7c,e). *Gliomastix* (Sordariomycetes)*, Suplenodomus* (Dothideomycetes)*, Knufia* <u>(Ascomycetes familia incertae)</u>*, Articulospora* (Leotiomycetes)*, and Ganoderma* (Agaricomycetes) accounted for 78% of the increase in group relative abundance of Winter-Adapted fungi from September to March in the Floodplain (Figure 7b). *Thelephora* (Agaricomycetes)*, Hebeloma* (Agaricomycetes)*, Archaeorhizomyces* (Archaeorhizomycetes), and *Tetracladium* (Leotiomycetes) were Spring-Adapted fungi with substantial increases in relative abundance in the Floodplain after snowmelt (Figure 7f). Very few fungal OTUs employed the Snowmelt-Specialist life strategy in the Floodplain (Table 4), which was composed of only two fungal genera [i.e., *Minutisphaera* (Dothideomycetes) and *Endosporium* (Dothideomycetes); Figure 7d]. All fungal OTUs with life strategies assigned are listed in Supplemental Table 2.

**Figure 7.**
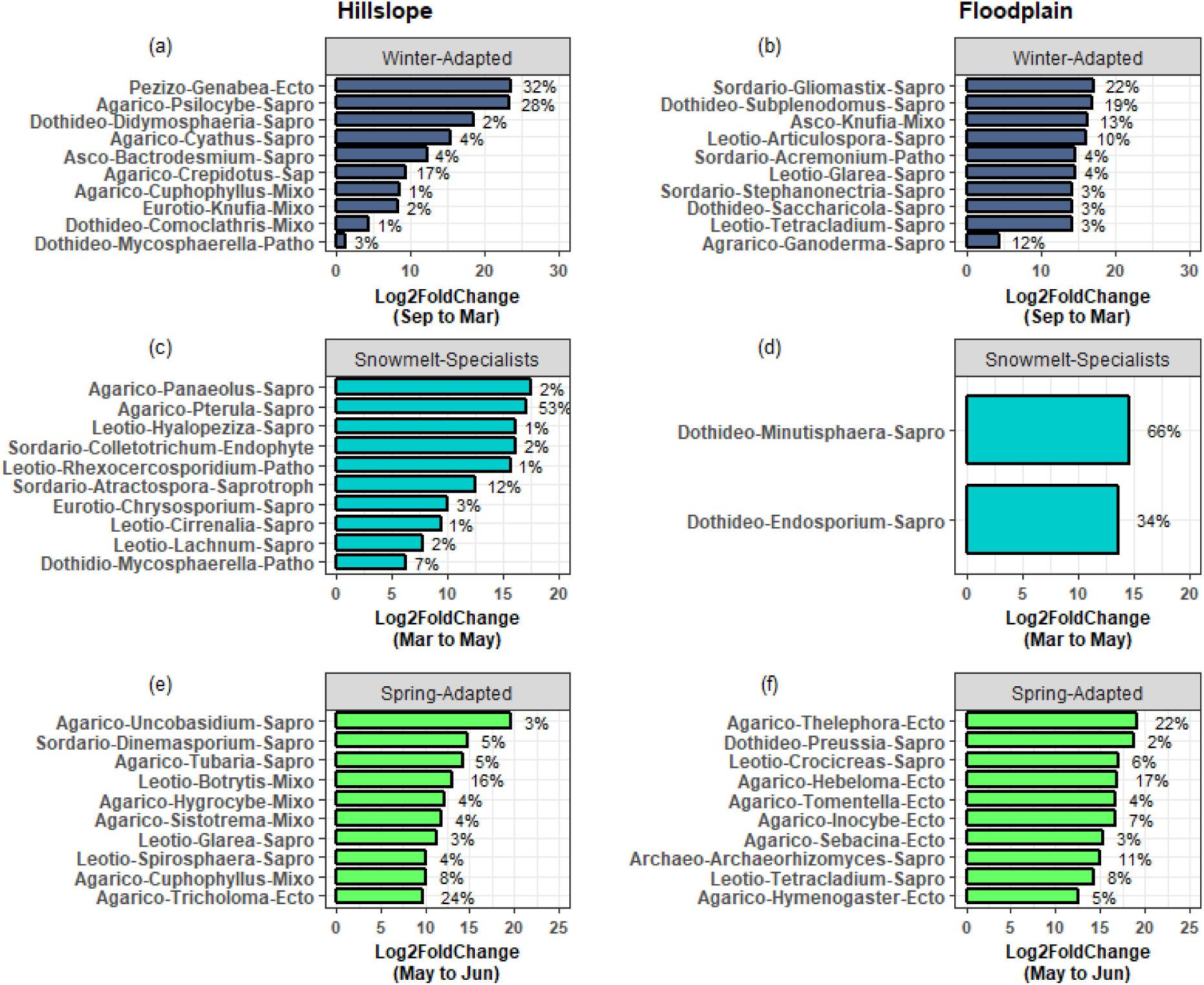
The top 10 fungal phylotypes within each life history strategy. Phylotypes were ranked by their percent change in abundance between the specified sampling time-points. The bars represent log2fold changes and the percent contributions to the change in group relative abundance are given at the side of the bar. Only two fungal genera contributed to the significant change in fungal snowmelt-specialists from March to May in the Floodplain location (panel d). Abbreviations for taxonomy (class-level) are Agarico – Agaricomycetes; Archaeo - Archaeorhizomycetes; Asco - Ascomycetes; Dothideo – Dothideomycetes; Eurotio – Eurotiomycetes; Leotio – Leotiomycetes; Pezizo – Pezizomycetes; Sordario - Sordariomycetes . Abbreviations for functional groups are Ecto – ectomycorrhizae; Endophyte – root endophyte; Mixo – mixotrophic; Patho – pathotrophic; Sapro - saprotrophic

## Discussion

Soil microbial biomass is commonly observed to bloom beneath the winter snowpack and subsequently decline following snowmelt, often resulting in a significant pulse of soil N in spring (Grogan and Jonasson 2003, Edwards et al. 2006, Schmidt et al. 2007). In this study, we primarily sought to identify bacteria, archaea, and fungi that were associated with the microbial biomass bloom during winter and biomass crash following snowmelt. We also sought to infer whether the traits that govern microbial community assembly during and after snowmelt were phylogenetically conserved. This was accomplished by measuring the phylogenetic relatedness of bacteria and archaea, or by comparing guilds of soil fungi, that were grouped into life strategies based upon relative abundance patterns before, during, and after snowmelt.

Overall, we found that (1) increases in soil microbial biomass production beneath the winter snowpack were observed at both locations in our study (Hillslope and Floodplain) and appeared to be triggered by a pulse of snowmelt infiltration (Figure 2), (2) the microbial biomass collapse was associated with a significant pulse of N measured as extractable soil NO3^-^ (Figure 2), (3) three microbial life strategies (Winter-Adapted, Snowmelt-Specialist, and Spring-Adapted) were identified at each soil depth at both locations (Figures 4, 6), (4) life strategies were most often phylogenetically clustered (bacteria and archaea) or shared similar trophic modes (e.g., saprotrophic fungi; Tables 3, 4), and that (5) a few select taxa within each life strategy contributed disproportionately to the abundance responses (Figures 5, 7). Thus, we have shown that bacteria, archaea, and fungi with similar responses to snow accumulation and snowmelt are likely to share adaptive traits and we conclude that this framework may be useful in understanding an organism’s snowmelt niche and response to changing winter snowpack conditions.

### Role of moisture and substrate utilization in structuring the Winter-Adapted niche

Phylogenetic clustering of Winter-Adapted archaea and bacteria may partly be explained by poor connectivity of soil pore spaces due to dry soil conditions that also lead to specific substrate utilization patterns during the winter. Because plant detritus is thought to be the primary substrate utilized by bacteria and fungi during winter (Taylor and Jones 1990, Uchida et al. 2005, Hooker and Stark 2012, Isobe et al. 2018), low substrate diffusion rates, combined with high spatial heterogeneity and low resource availability probably selects for bacteria and fungi with specialized abilities to degrade complex soil organic matter and plant root or leaf litter. A predominance of α-Proteobacteria (e.g., *Tardiphaga*, *Massilia*, *Phenylobacteria*, *Rhodoplanes*) known to be early-to mid-stage colonizers of decomposing root and leaf-litter (Purahong et al. 2016, Bonanomi et al. 2019) supports this hypothesis (Figure 5). Similar to the bacterial community, we observed that Winter-Adapted fungi were predominantly saprotrophic, with a few select taxa that are known wood or detritus decomposers (e.g., *Gliomastix*, *Psilocybe*, *Ganoderma*) contributing significantly to the increase in Winter-Adapted fungi (Figure 7). Additional fungal taxa commonly found during early-to mid-stages of decomposition of leaf or root litter (e.g., *Acremonium*, *Mycosphaerella*, *Saccharicola*, *Tetracladium*; Kuhnert et al. 2012) also had significant positive responses to winter snow accumulation at our study site.

Lipson et al. (2009) previously found that bacteria and fungi isolated from beneath the snowpack in winter had a propensity for high growth rates, high mass-specific respiration, and low growth yields. If those observations are generalizable to the winter-adapted taxa observed here, we hypothesize that such physiological traits would select for a winter-adapted phenotype characterized by enhanced investment in extracellular enzyme production to optimize nutrient acquisition *via* depolymerization of soil organic matter (Schimel et al. 2007, Malik et al. 2018, Zhalnina et al. 2018).

We also observed two potentially N-fixing genera (e.g., *Rhizobium* and *Bradyrhizobium*) that contributed significantly to the overwinter increase in Winter-Adapted archaea and bacteria on the Hillslope, but these genera were not observed in the Floodplain (Figure 5). Recent studies have shown high abundance of N-fixing *Bradyrhizobium* and *Rhizobium* in mid-to late-stages of litter decomposition and their presence is thought to alleviate N-limitation that occurs during late stages of decomposition, which thereby facilitates the activity of decomposers that degrade more complex biopolymers (Purahong et al. 2016, Lladó et al. 2017, Bonanomi et al. 2019). In seasonally snow-covered ecosystems, like the East River study site, a similar effect might occur as winter progresses due to the prolonged absence of fresh C inputs from plants aboveground and the potential for progressive N-limitation as the decomposer community propagates. In contrast to the Hillslope, we observed greater total soil N availability in the Floodplain as well as autotrophic, nitrifying archaea (e.g., Thaumarchaeota) during winter. These observations, along with a lack of a significant contribution from N-fixing genera (e.g., *Rhizobium* and *Bradyrhizobium*), might indicate that the N-fixing niche is not present beneath the snowpack in the Floodplain.

### Snowmelt infiltration triggers microbial biomass bloom

The microbial biomass bloom coincided with a period of extended soil saturation that occurred as a result of snowmelt infiltration and lasted for a period of approximately 65 days (Figures 1, 2). Snowmelt-triggered biomass production stands in contrast to previous hypotheses predicting that a sudden decrease in osmolarity due to snowmelt infiltration should cause lysis of microbial cells (Schimel et al. 2007, Jefferies et al. 2010). However, our results are similar to those of a study conducted in a sub-arctic hedge ecosystem that found a soil microbial biomass bloom initiated 45 to 60 days prior to soil becoming snow-free and occurring when soil temperatures were above 0 °C (Edwards et al. 2006).

We hypothesize that alleviation from moisture and substrate limitation may have triggered the microbial biomass bloom that we observed under the winter snowpack. While microbial metabolic activity has been shown to occur in soils with temperatures well below 0 °C (McMahon et al. 2009), our findings suggest that soil and air temperatures need to be warm enough to induce net snowmelt infiltration and for soil water to be in the liquid phase in order to trigger the soil microbial bloom. In addition, the microbial bloom did not occur in Floodplain soils deeper than 5 cm, where higher soil moisture availability existed throughout the winter.

We also observed phylogenetic overdispersion of Snowmelt-Specialist bacteria and archaea on the Hillslope in our study (Table 3), whereas Snowmelt-Specialists were phylogenetically clustered in the Floodplain. For the Hillslope, these results imply that ecologically dissimilar organisms with a broad phylogenetic distribution contribute to the biomass bloom during snowmelt and biomass collapse thereafter. Thus, antagonistic biotic interactions, such as competitive exclusion and density-dependent regulation of population size, are likely to be the community assembly mechanisms underlying the microbial biomass bloom, biomass crash, and taxonomic succession that we observed before and after snowmelt. For example, antagonism resulting from toxin production may promote biomass production by reducing competition during a time of high resource availability (Moons et al. 2009, Hibbing et al. 2010). After snowmelt, the population decline on the Hillslope also indicates far greater mortality compared to reproduction rates, which may result from resource depletion or heavy predation by viruses, microfauna, and mesofauna (Moons et al. 2009, Phaiboun et al. 2015, Georgiou et al. 2017).

Alternatively, competitive exclusion of Winter-Adapted organisms may occur during snowmelt if Snowmelt-Specialist archaea and bacteria have higher growth efficiencies compared to Winter-Adapted taxa (Georgiou et al. 2017). Dissolved organic substrate quality and quantity is known to increase during snowmelt (Campbell et al. 2014, Patel et al. 2018), thus organisms that are characteristic of the biomass bloom (i.e. Snowmelt-Specialists) should be able to re-allocate energy resources away from extracellular enzyme production and nutrient uptake toward protein/fatty acid/DNA synthesis to maximize growth efficiency (Schimel et al. 2007, Malik et al. 2018). Furthermore, it is unlikely that organisms with high growth efficiencies or whole communities with high rates of biomass production would be closely related phylogenetically, but rather could possess genomic features and adaptive traits that are shared due to convergent evolutionary trajectories (Roller et al. 2016, Muscarella and Lennon 2018).

### Spring Nitrogen Dynamics and Spring-Adapted Bacteria and Fungi

We attribute the pulse of extractable soil NO3^-^ observed in spring to elevated N mineralization and nitrification rates resulting from a flush of labile C and N that was released from microbial biomass after snowmelt. Lipson et al. (1999) similarly showed a pulse of soil protein after snowmelt that was similarly attributed to the lysis of soil microbial biomass. Furthermore, four ammonia or nitrite-oxidizing groups [e.g., 0319-6A21 (Nitrospirae), unassigned *Nitrospirae* spp., and unassigned *Thaumarchaeota* spp.] contributed substantially to the increase of Spring-Adapted archaea and bacteria on the Hillslope (Figure 5), which also coincided with the pulse of soil NO3^-^ in spring (Figure 2), suggesting substantial autotrophic nitrification. In contrast to the Hillslope, we did not observe NO3^-^ accumulation in the Floodplain despite a dramatic microbial biomass collapse in the upper soil profile and increases in the abundance of some nitrifying organisms. Lower nitrate accumulation could arise from any combination of the following: lower mineralization and nitrification rates, higher N uptake rates (plant and microbial) with higher N immobilization, or higher rates of NO3^-^ assimilation and denitrification at the Floodplain compared to the Hillslope.

In addition to the prevalence of ammonia-oxidizing genera, we also found that four groups of Acidobacteria (i.e., Subgroup 1, Subgroup 2, *Candidatus* Solibacter, and RB41) made large contributions to the increase in Spring-Adapted bacteria on the Hillslope (Figure 5e). Acidobacteria are traditionally considered to be oligotrophs with abundances negatively correlated to soil organic C (Fierer et al. 2007). However, an increasing number of studies have shown that Acidobacteria responses to gradients in organic C availability can be either positive or negative, challenging their status as strictly oligotrophic (Kielak et al. 2016, Lladó et al. 2017). In addition, some Acidobacteria can be isolated from the environment under anoxic conditions using microbial necromass (i.e., gellan gum) as a growth substrate and *Ca.*Solibacter has 4-to 6-fold higher nutrient-transporting genes in its genome compared to most other bacteria (Kielak et al. 2016). Therefore, some Acidobacteria phylotypes may occupy the Spring-Adapted niche by capitalizing on the necromass from the biomass crash along with high investment in nutrient transporters after snowmelt.

Mycorrhizal fungi made significant contributions to the increase in Spring-Adapted fungi in both the Hillslope and Floodplain (Figure 7); this pattern was driven primarily by an increase in ectomycorrhizal fungi. For example, in the Floodplain, 6 of the top 10 Spring-Adapted fungi were ectomycorrhizae. If ecosystem N retention in spring is enhanced by well-coupled plant-mycorrhizae phenology, then the implications of our results are noteworthy with respect to changing winter conditions. For example, although earlier snowmelt could result in an earlier pulse of N released from the crash in microbial biomass, floodplain ecosystems may buffer ecosystem N losses *via* well-coupled plant-mycorrhizal N uptake in spring. It may also be possible to predict the capacity of other watershed locations to retain N in response to earlier snowmelt based upon plant distributions and plant-mycorrhizae associations, given that plant species assemblages and plant-mycorrhizal associations can be mapped and inferred at high spatial resolution using remote-sensing methods (Fisher et al. 2016, Falco et al. 2019).

### Conclusions

Because snowmelt rates have been declining in mountainous regions in recent decades (Musselman et al. 2017, Harpold and Brooks 2018), the length of time during which soils are saturated beneath a melting snowpack may increase in the future and thereby promote a larger snowmelt microbial biomass bloom and crash. Alternatively, a future with warmer air temperatures and a shallower winter snowpack would be expected to result in an increase in the frequency of sustained soil freezing or more frequent soil freeze-thaw events during winter (Hardy et al. 2001, Brown and DeGaetano 2011, Kreyling and Henry 2011), which would most likely reduce the magnitude of the microbial biomass bloom, biomass collapse, and pulse of N after snowmelt (Brooks et al. 1998, Buckeridge et al. 2010, Ueda et al. 2013). Whether future environmental conditions will sustain an overwinter microbial bloom warrants further study, because the nutrient flush from microbial biomass following snowmelt can be one of the largest annual soil N pulses in some high-latitude or high-altitude ecosystems (Grogan and Jonasson 2003, Schmidt et al. 2007).

Here, we have provided novel evidence that the snowmelt period is an environmental filter that differentiates soil bacteria, archaea, and fungi into three distinct life strategies. The phylogenetic relatedness of bacteria and archaea and enrichment of fungal functional groups within life strategies suggests that selective, deterministic processes allow Winter-Adapted, Snowmelt-Specialist, and Spring-Adapted organisms to occupy distinct niches that are related to the winter snowpack and snowmelt. Although our main goal was not to compare and contrast two distinct watershed locations (Hillslope and Floodplain), we did observe differences in the magnitude of the microbial biomass bloom and collapse among locations and soil depths, patterns of NO3^-^ accumulation in spring, and differences in contributions from various taxa to each life strategy. These results indicate that local factors, in part, shape the response of these watershed locations to snowmelt. We contend, however, that common microbial trait distributions allow for ecologically similar organisms to occupy the same life strategy, in spite of watershed location. This hypothesis can be tested using high-resolution, culture-independent methods (e.g., genome-resolved metagenomics and metatranscriptomics) and by quantifying traits in laboratory isolates to further explore the microbial snowmelt niche.

## Acknowledgements

We would like to thank Erica Dorr, Biz Whitney, Hans Wu Singh, and Chelsea Wilmer for their assistance in the lab and in the field. Mark Conrad, Rosemary Carroll, and Wendy Brown were extremely helpful digging snow pits during winter. We also wish to thank Jenny Reithel and Shannon Sprott at Rocky Mountain Biological Laboratory (RMBL) for their help at the field site. RMBL lab spaces and equipment are funded in part by the RMBL Equipment (Understanding Genetic Mechanisms) Grant, DBI-1315705. This material is based upon work supported as part of the Watershed Function Scientific Focus Area funded by the U.S. Department of Energy, Office of Science, Office of Biological and Environmental Research under Award Number DE-AC02-05CH11231.

## Author Contributions

POS, HRB, and ELB designed the study, which is a contribution to the Watershed Function Scientific Focus Area lead by SSH and KHW. Data collection and analysis was performed by POS (primarily) and SW, AP, MB, UK, NJB, HRB, and ELB. POS, HRB, and ELB wrote the manuscript and all authors contributed to revising the manuscript.

